# Mate selection provides similar genetic progress and average inbreeding than optimum contribution selection in the long-term

**DOI:** 10.1101/2020.01.13.904219

**Authors:** Grazyella Massako Yoshida, José Manuel Yáñez, Sandra Aidar de Queiroz, Roberto Carvalheiro

## Abstract

Optimum contribution selection (OCS) and mate selection (MS) are alternative strategies to maximize genetic gain under controlled rates of inbreeding. There is evidence in the literature that MS outperforms OCS in controlling inbreeding under the same expected genetic gain in the short-term. It is unclear, however, if the same would occur in the long-term. This study aimed to compare OCS and MS regarding short and long-term genetic progress and inbreeding, using simulated data. The structure of the simulated population aimed to mimic an aquaculture breeding program. Twenty discrete generations were simulated, considering 50 families and 2,000 offspring per generation, and a trait with a heritability of 0.3. OCS and MS were applied using a differential evolution (DE) algorithm, under an objective function that accounted for genetic merit, inbreeding of the future progeny and coancestry among selection candidates. For OCS, the optimization process consisted of selection based on optimum contribution followed by minimum inbreeding mating. Objective functions using different weights on coancestry were tested. For each application, 20 replicates were simulated and the results were compared based on their average. Both strategies, OCS and MS, were very effective in controlling inbreeding over the generations. In the short-term, MS was more efficient than OCS in controlling inbreeding under the same genetic gain. In the long-term, OCS and MS resulted in similar genetic progress and average inbreeding, under the same penalty on coancestry.

## 1. Introduction

Selective breeding schemes for animals involve selection and mating decisions, i.e. the definition of the animals to be used as parents of the next generation and the decision regarding the mates to be performed among the selected parents. Both decisions have a high impact on the outcome of the breeding programs because they determine genetic gain and inbreeding level of subsequent generations (Falconer and Mackay, 1996).

Aquaculture breeding is usually characterized by having controlled populations with limited size and high fecundity. The latter allows applying high selection intensity, which enables relatively high annual genetic gain in the short-term. However, this can also potentially lead to very few families dominating the genetic contributions to the next generation and, consequently, the rate of inbreeding can easily become increased to undesirable levels (Caballero et al., 1996). High rates of inbreeding may have an important effect on medium- and long-term response to the selection, increase the manifestation of deleterious alleles, cause inbreeding depression, reduce the genetic variation and the genetic progress of subsequent generations (Falconer and Mackay, 1996).

Different strategies have been proposed to control inbreeding such as optimum contribution selection (OCS) (Meuwissen, 1997; Woolliams and Thompson, 1994), which maximizes genetic response while constraining inbreeding by restricting the coancestry among the selected parents. The OCS is usually applied with the combination of mating strategies as minimum inbreeding and compensatory mating and has been proved to be effective in controlling the long-term inbreeding (Grundy et al., 1998; Meuwissen, 1997; Sonesson and Meuwissen, 2000).

An alternative strategy is to perform selection and mate allocations simultaneously by using mate selection (MS) (Kinghorn and Shepherd, 1999; Shepherd and Kinghorn, 1999). In this approach, different components related to the breeding objectives can be accommodated in an objective function (OF) that, when optimized, results in a mating list that indirectly determines the contribution of each selection candidate (Kinghorn, 2011). The main challenge is to determine a proper OF and a method to optimize it (Kinghorn and Shepherd, 1999). MS has also been proved to be effective in controlling inbreeding while maximizing genetic gain, and other components of the objective function when present (Carvalheiro et al., 2010a; Hayes et al., 2002; Kinghorn and Shepherd, 1999; Kinghorn, 2011, 1998; Li et al., 2006; Shepherd and Kinghorn, 1999).

Using real Nile tilapia and coho salmon datasets, Yoshida et al. (2017) observed that, under the same expected genetic gain, MS outperformed OCS in controlling inbreeding in the short-term. However, their data did not allow contrasting both strategies in the long-term and a simulation study was recommended. The objective of the present study was to compare OCS and MS in aquaculture breeding, regarding short and long-term genetic progress and inbreeding, using simulated data. Partial results of this study were presented by Yoshida et al. (2018a).

## 2. Material and Methods

### 2.1 Simulated data

The simulation process aimed to mimic the structure of a fish population. The animals from the base population (G0) comprised 25 males and 50 females. Their additive genetic values (g_i_) were sampled from a Gaussian distribution 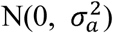, where 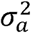 is the additive genetic variance 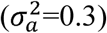 in G0. Phenotypes in the base population were then calculated as:

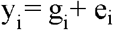

where e_i_ is the environmental effect sampled from 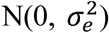, with 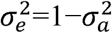. Thus, in the base generation, the phenotypic variance was equal to 1 and the heritability equal to 0.3. Phenotypic values of the offspring in later generations were simulated as:

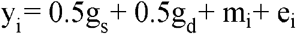

where g_s_ and g_d_ are the additive genetic values of the sire and dam, respectively; m_i_ is the Mendelian sampling effect with the distribution 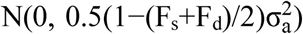, with F_s_ and F_d_ being the inbreeding coefficients of the sire and dam, respectively; and e_i_ was defined as above. Each female produced 40 offspring with equal probability of being a male or a female, totaling 2,000 progenies per generation. For each scenario encompassing a selection and mating strategy, 20 discrete generations were simulated (G1-G20).

### 2.2 Objective function

An algorithm based on Differential Evolution (DE) (Storn and Price, 1997) was developed in FORTRAN language by Carvalheiro et al. (2010b), which allows applying either OCS or MS depending on the weights used for the different components of the objective function (OF) to be optimized. The OF used was:

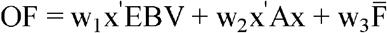

where, x’EBV is the expected merit of the future progeny; x’Ax is the weighted mean coancestry of selected parents; 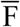 is the expected average inbreeding coefficient of the future progeny; w_1_ to w_3_ are the corresponding weighting factors and x is the vector to be optimized of genetic contributions for each candidate (the symbol ’ denotes a transposed vector).

Although not explicitly described in the OF, the mate allocations were determined by 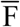, following the problem representation suggested by Gondro and Kinghorn (2008). In this representation, an auxiliary vector is used internally in the mate selection algorithm with the number of elements equal to the number of mates. Each element of the auxiliary vector is a real number used to indirectly determine the mates. These real numbers are ranked, and the resultant rankings ultimately define the mates to be performed (Yoshida et al., 2017).

For each discrete generation, it was considered as selection and mate candidates the best female (50 animals) and the best five males (250 animals) per family. The males were allowed to be mated with a maximum of four females and each female was mated once, i.e. the contribution of each female was not optimized.

The operational parameters of the DE algorithm to optimize the OF where: population size = 2 times the number of candidates; crossover rate = 0.5; mutation factor = 0.2 (or 0.9 every 4 generations); and maximum number of generations of the evolutionary process (maxgen = 100,000). Convergence was assumed when the range and the mean absolute deviation of the OF, considering all the possible solutions per generation, were lower than 1×10^−6^. The best solution from the maxgen generation was considered as the optimum solution when the convergence criterion was not attained.

The approach proposed by Lampinen and Zelinka (1999) was adopted to provide integer solutions for the number of mates per candidate. To increase computational efficiency, Colleau (2002) indirect approach was adopted to calculate coancestry, and linked lists (Knuth, 1975) were used for the storage and calculations involving sparse matrices. The mate selection algorithm is freely available for research purpose and can be obtained under request to the corresponding author.

### 2.3 Scenarios

Initially, four different OF were optimized, characterizing two applications of OCS (OCS1 and OCS2) and two of MS (MS1 and MS2). They differed according to the weights used for coancestry (*w*_*2*_) and inbreeding (*w*_*3*_). OCS1 and MS1 used a *w*_*2*_ of −10. A higher emphasis on coancestry was given on OCS2 and MS2 (*w*_*2*_ = −20). For OCS1 and OCS2, a *w*_*3*_ equal to −0.00001 was used, corresponding to the application of OCS followed by minimum inbreeding mating. For MS1 and MS2, a *w*_*3*_ equal to −1 was used, so in these two OF the selection and mating were performed simultaneously. All OF used the same weight for the expected merit of the future progeny (*w*_*1*_ = 1). In summary, the following weights were used for each application: OCS1 (*w*_*1*_=1, *w*_*2*_=-10, *w*_*3*_=-0.00001); OCS2 (*w*_*1*_=1, *w*_*2*_=-20, *w*_*3*_=-0.00001); MS1 (*w*_*1*_=1, *w*_*2*_=-10, *w*_*3*_=-1) and MS2 (*w*_*1*_=1, *w*_*2*_=-20, *w*_*3*_=-1). These weights were determined empirically based on results from previous studies (Carvalheiro et al., 2010a; Yoshida et al., 2017).

In addition, three other OF were applied for comparison purpose. They were: truncation selection (TS) followed by random mating (TS1: *w*_*1*_=1, *w*_*2*_=0, *w*_*3*_=0); truncation selection followed by minimum inbreeding mating (TS2: *w*_*1*_=1, *w*_*2*_=0, *w*_*3*_=-0.00001) and OCS followed by random mating (OCS3: *w*_*1*_=1, *w*_*2*_=-10, *w*_*3*_=0). For each scenario, 20 replicates per generation (representing different populations) were simulated and their results were averaged for the comparisons. The results were compared based on genetic gain, coancestry, and inbreeding, provided by the optimization of each OF.

## 2. Results

OCS and MS were applied using a differential evolution algorithm, under an objective function that accounted for genetic merit, inbreeding of the future progeny and coancestry among selection candidates. For OCS, the optimization process consisted of selection based on optimum contribution followed by minimum inbreeding mating (OCS1 and OCS2) or random mating (OCS3). Objective functions were tested with a lower (OCS1, OCS3 and MS1) or higher (OCS2 and MS2) emphasis on constraining coancestry. Truncation selection (TS) followed by random mating (TS1) and by minimum inbreeding mating (TS2) was also applied for comparison.

Figure 1 presents the genetic response of the different strategies along the generations. OCS1 presented a genetic gain equivalent to MS1 and the genetic gain of OCS2 was similar to MS2, in the short and long term. Therefore, the contrasts between OCS1 *vs* MS1 and OCS2 *vs* MS2 allow reasonable comparisons between both strategies in terms of controlling inbreeding under the same rate of genetic gain (discussed later). As expected, increased penalty on coancestry resulted in the reduction of genetic response. For instance, at G20, OCS2 presented an average breeding value (7.45) 7.22% lower than OCS1 (8.03). Truncation selection strategies (TS1 and TS2) presented the highest genetic gain, which was about 11-12% superior to those presented by OCS1 and MS1 at G20. OCS followed by random mating (OCS3) presented similar genetic response than OCS followed by minimum inbreeding mating (OCS1) (Figure 1).

**Figure 1.**
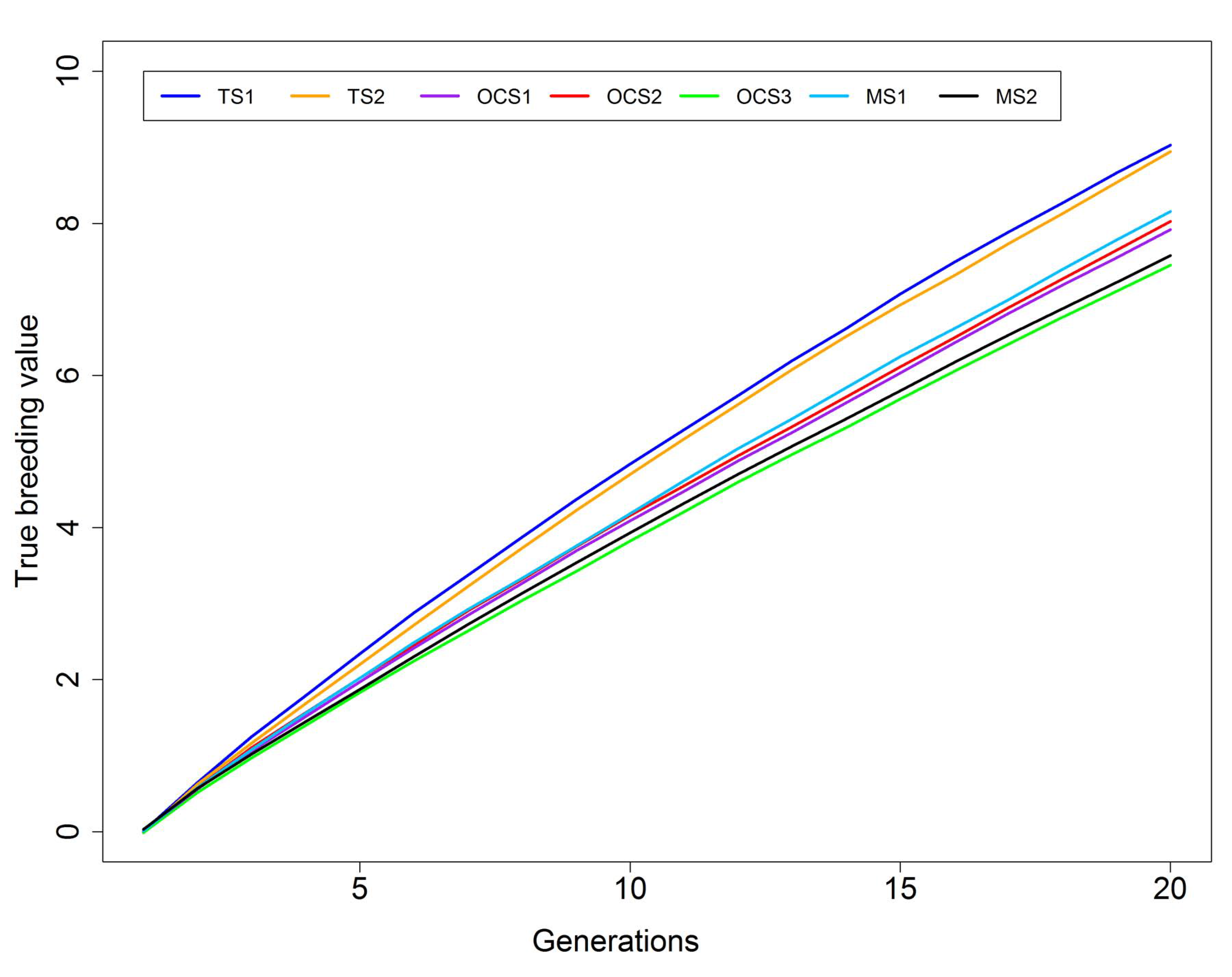
Average true breeding value over 20 generations of selection considering different objective functions. TS1: truncation selection (TS) followed by random mating (RM); TS2: TS followed by minimum inbreeding mating (MIM); OCS1: optimum contribution selection (OCS) followed by MIM, with lower emphasis on coancestry; OCS2: OCS followed by MIM, with higher emphasis on coancestry; OCS3: OCS followed by RM, with lower emphasis on coancestry; MS1: mate selection (MS) with lower emphasis on coancestry; MS2: MS with higher emphasis on coancestry.

The variance of true breeding values decreased in the first five generations and kept almost constant in the next generations for the different strategies except for truncation selection (TS1 and TS2), which showed a continuing decrease of genetic variance along the generations. At G20, the genetic variance was around 0.23-0.24 for the OCS and MS strategies, whereas it was close to 0.14-0.15 for the TS strategies (Figure 2).

**Figure 2.**
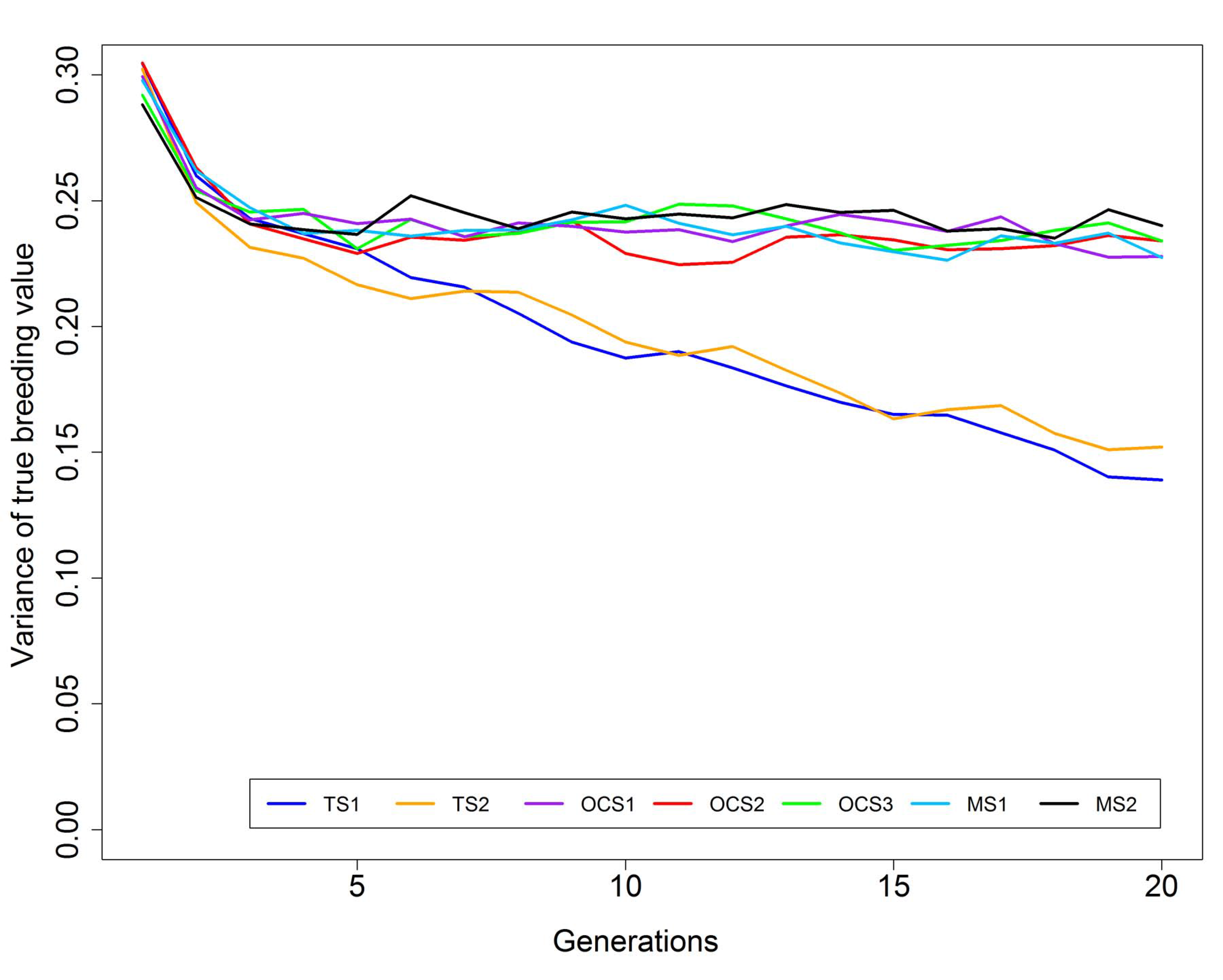
Variance of true breeding value over 20 generations of selection considering different objective functions. The scenarios definition is present in Figure 1.

Similar values of coancestry were observed between OCS1 and MS1 and between OCS2 and MS2, increasing up to 0.21 for OCS1 and MS1 and up to 0.16 for OCS2 and MS2 at G20. TS strategy resulted in a much higher level of coancestry than OCS or MS. At G20, coancestry of TS1 and TS2 were equal to 0.94 and 0.89, respectively. OCS1 and OCS3 presented similar coancestry trends (Figure 3).

**Figure 3.**
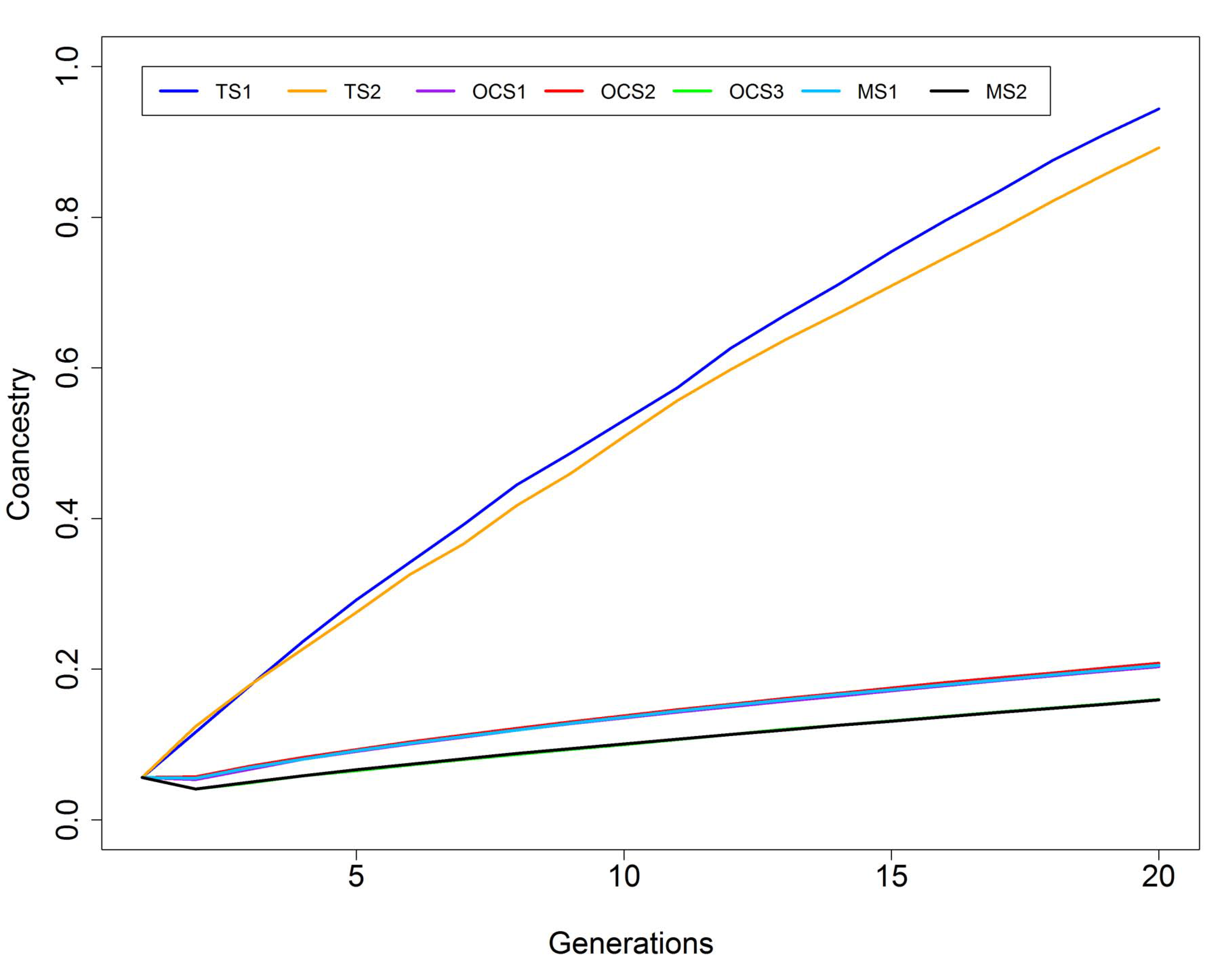
Average coancestry over 20 generations of selection considering different objective functions. The scenarios definition is present in Figure 1.

The number of sires selected by generation varied among the strategies (Figure 4). For TS1 and TS2, the number of sires selected was constant and equal to 13 along the generations. For OCS and MS, the number of sires selected increased along the generations reaching a plateau that corresponded to the maximum possible value, i.e. 50 sires selected to be mated with 50 females. OCS1 and MS1 started using less sires (27) than OCS2 (37) and MS2 (38), but also ended up using 50 sires in the last simulated generation (Figure 4).

**Figure 4.**
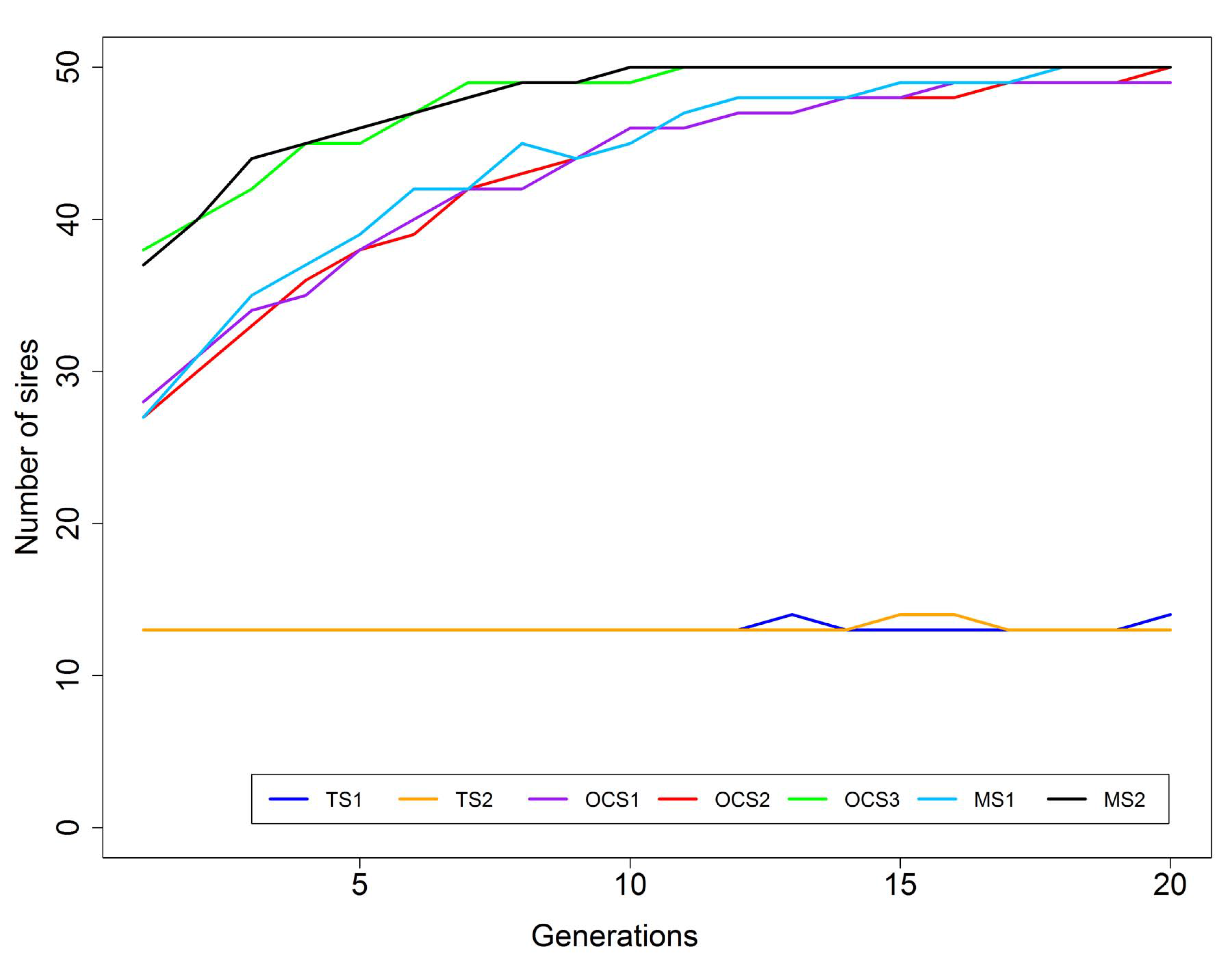
Average number of sires selected over 20 generations of selection considering different objective functions. The scenarios definition is present in Figure 1.

In the short-term, MS was more efficient in controlling inbreeding than OCS under the same penalty on coancestry (Figure 5A). At G4, for example, the average inbreeding of the different strategies was equal to 1.61% and 0.88% for OCS1 and MS1, and equal to 0.89% and 0.36% for OCS2 and MS2, respectively. However, in the long-term, MS and OCS presented similar inbreeding level under the same penalty on coancestry (Figure 5B). For instance, the average inbreeding of OCS1 and MS1 at G20 was equal to 8.78% and 8.49%, respectively. MS and OCS, which accounted for coancestry, were very efficient in controlling inbreeding compared to truncation selection followed by mating minimizing inbreeding (TS2), which resulted in an average inbreeding of 41.06% at G20. This value was even higher for truncation selection followed by random mating (44.37%). OCS3 presented higher inbreeding rate in the short-term compared to the other OCS and MS strategies (Figure 5A). After G5, OCS3 presented an inbreeding rate similar to OCS1 and MS1, besides presenting an inbreeding level slightly higher (Figure 5B).

**Figure 5.**
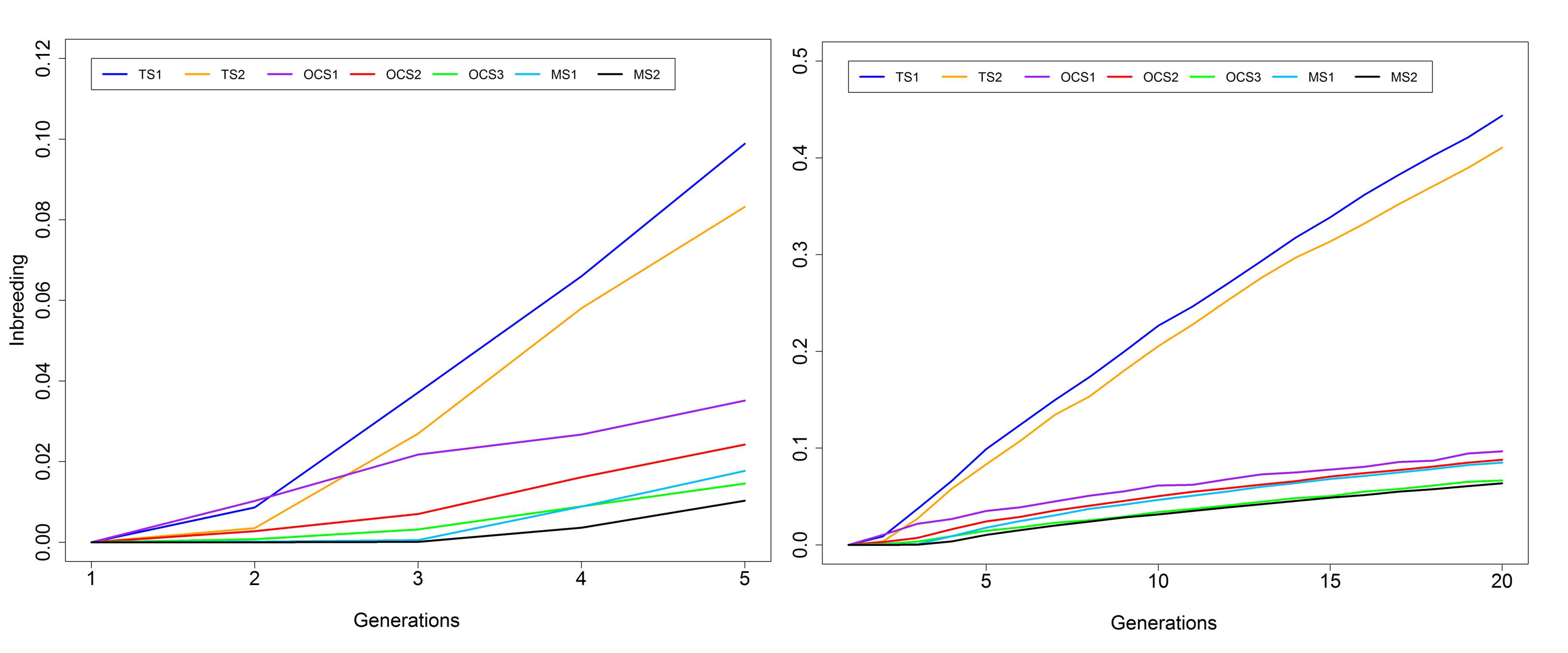
Average inbreeding level over 5 generations (a) and over 20 generations (b) of selection considering different objective functions. The scenarios definition is present in Figure 1. Note that the scale is not the same for Figure 5a and 5b.

## 3. Discussion

OCS is usually compared with TS regarding their genetic response under the same rate of inbreeding (Meuwissen and Sonesson, 1998). In our simulation, TS was not enforced to present the same rate of inbreeding than the other strategies. In contrast, TS was applied to focus on maximizing the genetic response of the next generation, with a constrain on the maximum allowed number of mates per candidate sire, which was equal to four for all strategies. As a result, TS presented the highest genetic response for having the highest selection intensity among the tested strategies. TS would probably not have the highest genetic response if inbreeding depression and deleterious recessive alleles were considered in the simulation process. It has been estimated that inbreeding depression corresponds, on average, to a decrease of about 0.137% of the mean of a trait per 1% of inbreeding (Leroy, 2014). In our simulated populations, this figure would correspond to reducing the mean up to 6% for TS (F>40%) and 1% or less for the remaining strategies (F<10%).

As reported in the literature (e.g. Woolliams et al., 2015), the strategy of performing minimum inbreeding mating after truncation selection (TS2) was not effective in controlling inbreeding. The only effective way to control inbreeding, as shown by OCS and MS results, is by penalizing or constraining the coancestry among selected parents, which is in accordance with the OCS theory (Meuwissen, 1997; Woolliams and Thompson, 1994). The penalties applied on coancestry here, allowed OCS and MS to present a substantial reduction on inbreeding compared to TS (up to 80-85%) and the decrease on the genetic gain, for selecting more sires and presenting a lower selection intensity, was not so expressive (up to 10-15%). Meuwissen and Sonesson (1998) observed that, at the same rate of inbreeding, OCS attained up to 44% more genetic gain than TS. Our results are not directly compared with theirs because, among other differences in the simulation process, we did not enforce TS to present the same rate of inbreeding than OCS, as previously discussed.

Sonesson and Meuwissen (2000) observed that OCS combined with non-random mating did not reduce the rate of inbreeding in comparison with OCS and random mating, probably because their OCS implementation enforced a constant rate of inbreeding across generations. The authors also observed that non-random mating allowed higher selection intensity for OCS, resulting in an increased genetic gain compared to random mating. In our simulation, OCS followed by minimum inbreeding mating (OCS1) resulted in a lower level of inbreeding and a similar genetic response than OCS combined with random mating (OCS3). So, depending on the implementation, OCS combined with non-random mating can allow increased genetic gain under the same rate of inbreeding (Sonesson and Meuwissen, 2000) or lower inbreeding under similar genetic gain (our results), compared to OCS with random mating.

Accounting for selection and mate decisions simultaneously (MS), and not in two steps as in OCS, allowed a better control of inbreeding in the short-term, under the same response to selection. This result is in agreement with Yoshida et al. (2017), who also observed evidence that MS outperformed OCS in controlling inbreeding in the short-term, for a real Nile tilapia low inbred population. This is probably associated to the fact that in the initial generations, when most animals were unrelated, MS was able to identify candidate sires that would result in similar genetic gain and coancestry than OCS, but with a smaller inbreeding of the progeny. This, however, was not the case in the long-term, where MS and OCS presented similar genetic gain and inbreeding. Therefore, after a certain level of coancestry and inbreeding, it seems that MS and OCS would select the same candidates irrespective of considering inbreeding of the progeny in the selection process (MS) or as a second step in the mating strategy (OCS).

Giving more emphasis on coancestry (OCS1 and MS1 vs OCS2 and MS2) resulted in a better control of inbreeding along the generations, with a small reduction on the genetic response. Unfortunately, the optimal weights to be used for the different components of the OF cannot be analytically determined in the evolutionary algorithm implemented. They need to be determined empirically as stressed by Yoshida et al. (2017). To overcome this drawback, the authors recommended to run the algorithm several times varying the weights for the different components, explore the potential outcomes and choose the set of weights that would result in a better balance among the different components, according to the goal of the breeding program. The advantage of using this strategy is that there is no need to determine a prior target value for the rate of inbreeding, which is usually determined empirically in most OCS applications (Woolliams et al., 2015).

A potential advantage of MS over OCS not investigated in the present study is related to its flexibility in incorporating different components to be concomitantly optimized in the objective function, together with genetic gain and coancestry/inbreeding, such as non-additive effects for multi-breed populations (Hayes et al., 2002), connectedness among contemporary groups (Carvalheiro et al., 2010a), genetic variability of the progeny (Yoshida et al., 2018b), among others (Kinghorn and Shepherd, 1999).

## 4. Conclusions

In the short-term and under the same genetic gain, MS was more efficient in controlling inbreeding than OCS followed by minimum inbreeding mating. However, in the long-term, MS and OCS presented similar results regarding the genetic gain and level of inbreeding.

## Acknowledgments

GMY acknowledge Fundação de Amparo à Pesquisa do Estado de São Paulo (FAPESP processes numbers 2014/20626-4 and 2015/25232-7) for Doctoral fellowship. RC acknowledge Conselho Nacional de Desenvolvimento Científico e Tecnológico (CNPq, processes numbers 308636/2014-7 and 305435/2017-5) for fellowship.

## Author’s contributions

GMY performed the simulations, MS analysis and wrote the initial version of the manuscript. JMY and SAQ contributed with study design and discussion. RC developed the MS program, designed the study, supervised work of GMY and contributed to the analysis, discussion and writing. All authors read and approved the final manuscript.

## Availability of data and material

Only simulated data was used in this study. The datasets and MS program used during the current study are available from the corresponding author upon reasonable request.

